# Ca^2+^-calmodulin regulates Kv7.1 channel gating by allosterically interfering with its inactivation path

**DOI:** 10.1101/2025.09.02.673758

**Authors:** Nitzan Daus, Flavio Costa, Carlo Guardiani, Maya Lipinsky, Ariel Ben-Bassat, Yoni Haitin, Alberto Giacomello, Bernard Attali

## Abstract

Like in many voltage-gated K^+^ channels (Kv), inactivation of the cardiac Kv7.1 channel is voltage-dependent but does not exhibit the hallmarks of N-type or C-type mechanisms. This peculiar inactivation is observed in wild-type channels and is exacerbated in many Kv7.1 mutations triggering cardiac arrhythmias. Previously, we showed that Kv7.1 inactivation could strikingly be prevented by Ca^2+^-calmodulin (Ca^2+^-CaM). Thus, how can Ca^2+^-CaM, localized at the channel inner boundaries, prevent inactivation that occurs distantly at the outer pore region and converges to the selectivity filter? Here, using network analysis, molecular dynamics simulations, and electrophysiology, we identify the inactivation paths coupling the voltage sensor domain to the selectivity filter, involving helices S1 and S6, and the P-Helix, which represents the underlying mechanism of Kv7.1 inactivation. Moreover, our data reveal the allosteric coupling mechanisms by which Ca^2+^-CaM signals interfere with the inactivation paths and prevent channel inactivation.

## Introduction

Inactivation of most voltage-gated K^+^ channels (Kv) occurs by N-type or/and C-type mechanisms ^1, 2, 3, 4, 5^. The fast N-type inactivation involves the occlusion of the inner pore by an intracellular peptide corresponding to the N-terminus of channel α or β subunits ^5^, while the slower C-type inactivation is reflected by structural rearrangements in the outer pore and by a hallmark sensitivity to high external K^+^ due to interactions between the permeant ions and the selectivity filter (SF) ^4, 6^. Currently, the molecular mechanisms underlying filter conformational changes during C-type inactivation are still a matter of controversy, with studies suggesting either filter constriction and/or pore dilation ^7, 8, 9, 10, 11, 12^.

Like many Kv channels, the inactivation of the cardiac Kv7.1 (or KCNQ1) channels is voltage-dependent but does not exhibit the hallmarks of N-type or C-type mechanisms, notably by lacking sensitivity to high external K^+^ concentrations ^13, 14, 15, 16, 17^. In the heart, Kv7.1 associates with the auxiliary subunit KCNE1 to produce the I_Ks_ K^+^ current, critical for the repolarization of the cardiac action potential ^18, 19^. Numerous mutations in Kv7.1 (KCNQ1) and KCNE1 are linked to cardiac arrhythmias such as atrial fibrillation and long QT syndrome (LQTS), predisposing patients to sudden cardiac death. Remarkably, many LQTS mutants exhibit strong inactivation ^20, 21, 22^.

Kv7.1 belongs to the family of domain-swapped Kv7 (KCNQ) channels that comprises five members and presents the unique feature of an absolute requirement to associate with the lipid, phosphatidylinositol 4,5-bisphosphate (PIP2) and calmodulin (CaM) for proper voltage-dependent gating ^23, 24, 25, 26^. The canonical electromechanical coupling infers that during gating, the movement of the S4 helix leads to a translation of the linker S4-S5, which couples the voltage sensor domain (VSD) motion to conformational changes of the pore ^27, 28, 29^. Previous studies reported that Kv7.1 features a two-open-state gating mechanism, showing that its VSD can trigger pore opening when activated to both intermediate (I) and fully activated (A) states, leading to two conducting open states: the fast intermediate open (IO) and the slow fully activated open (AO) ^14, 30, 31, 32, 33, 34^. Previously, we found that these two open states (also, denoted O1 and O2 open states) relax, respectively, into two discrete inactivation states – a slow, voltage-dependent, Ca^2+^-sensitive inactivation state I1 and a fast, voltage-dependent, Ca^2+^-insensitive inactivation state I2 ^35^. Notably, in wild-type (WT) Kv7.1 channels, inactivation I1 can be robustly increased by the removal of external Ca^2+^ ^35^. We showed that E295 and D317, located in the turret region and pore entrance, respectively, are involved in Ca^2+^ coordination, allowing D317 to form hydrogen bonding with the pore helix (P-Helix) residue W304, stabilizing the SF and preventing inactivation ^35^. The inactivation I1 seen in WT Kv7.1 under external Ca^2+^-free conditions reflects an intrinsic inactivation gating that is detected recurrently in many Kv7.1 LQTS mutations in normal Ca^2+^-containing solutions ^20, 21, 22, 36^.

A unique feature of WT Kv7.1 inactivation I1 (the only one considered in this work) resides in its ability to be prevented by Ca2+-CaM ^35^. For this reason, an allosteric mechanism that couples Ca^2+^-CaM, localized at the intracellular boundaries of the channel, with the SF, which controls inactivation, is accounted for. Network analysis of Molecular Dynamics (MD) simulations has been widely used to identify long-range communication paths between distant regions in proteins, including both transmembrane proteins such as ion channels and soluble proteins like E2 enzyme or tRNA synthetases ^37, 38^. In this work we combined network-theoretical analysis with MD simulations and patch-clamp electrophysiology to reveal allosteric paths involved in Kv7.1 inactivation. First, we revealed inactivation paths coupling the VSD to the SF; then, we identified communication paths between Ca^2+^-CaM and SF, providing mechanistic insights into the allosteric modulation of Kv7.1 inactivation by Ca^2+^-CaM. Electrophysiology data validated the theoretical predictions, showing that mutations of residues, acting as hubs in the communication network, potently impact inactivation and its modulation by Ca^2+^-CaM.

## Results

### Inactivation paths

In the absence of an inactivated Kv7.1 structure, MD simulations were run starting from two cryo-EM structures solved by the MacKinnon’s lab ^39, 40^, KCNQ1-CaM (PDB: 6UZZ) and KCNQ1-KCNE3-CaM-PIP2 (PDB: 6V01) (Fig. 1a) that show fully activated VSD conformations, which is an important prerequisite since Kv7.1 inactivation is voltage dependent. As in our previous work ^41^, we studied inactivation paths based on equilibrium MD simulations of the open state. This is justified because 1) in the absence of an inactivated structure, the open state is the closest available structure, differing only by the conformation of the SF, 2) inactivation starts from the open channel and, thus, the equilibrium fluctuations around the open state are expected to reflect accurately the allosteric path, at least in its initial part.

**Figure 1:**
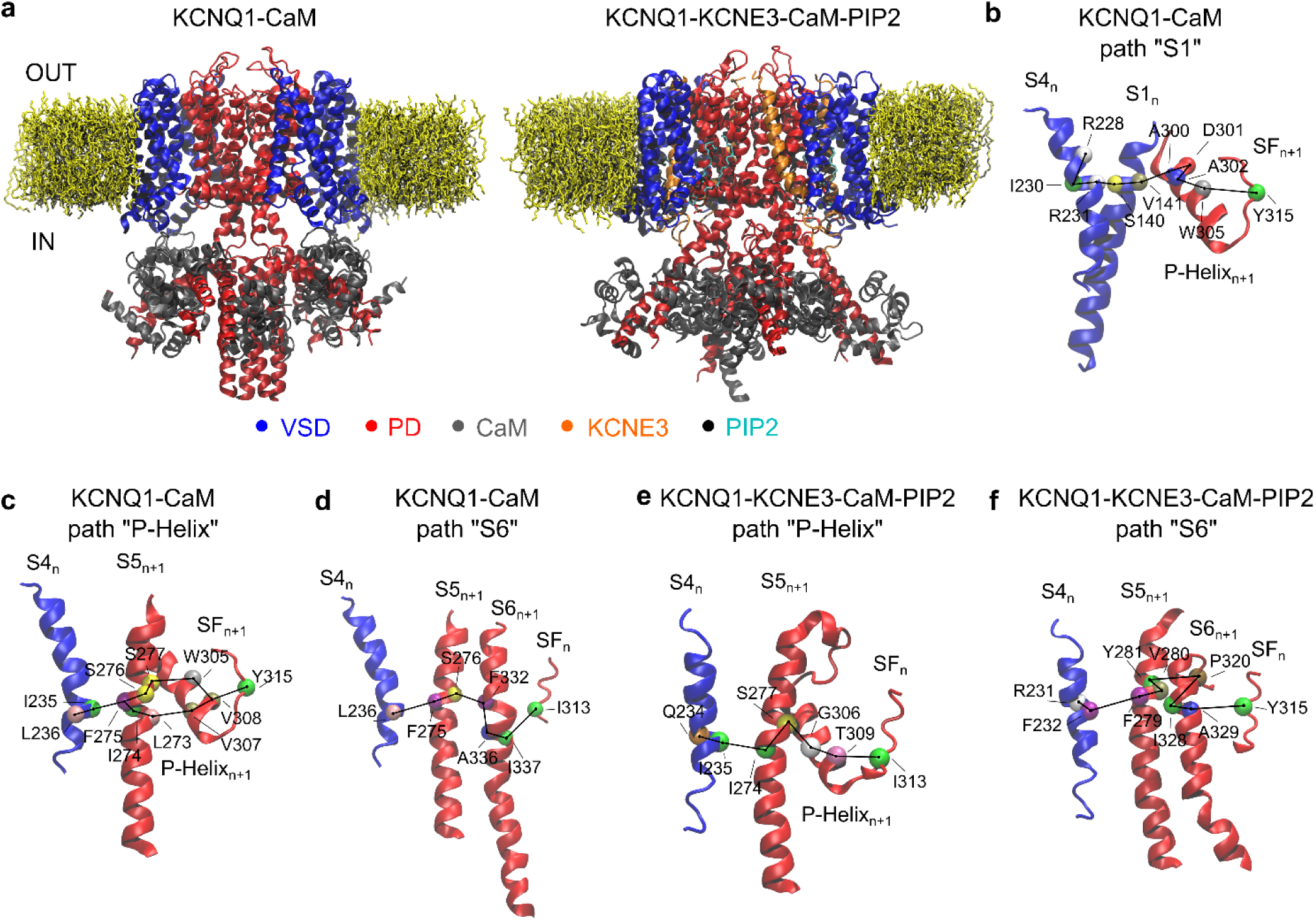
Inactivation paths predicted by the network analysis. **a** Side view of KCNQ1-CaM and KCNQ1-KCNE3-CaM-PIP2 channels embedded in lipid bilayers and coloured by domain: in blue, the voltage sensor domain ranging from helix S1 to S4-S5 loop; in red, the pore domain composed of helix S5, P-Helix, selectivity filter, and helix S6; in grey, the calmodulin; in orange, the KCNE3; in cyan, the phosphatidylinositol 4,5-bisphosphate (PIP2). Panels **b-d** show the inactivation paths “S1”, “P-Helix” and “S6” predicted in the KCNQ1-CaM system while panels **e-f** “P-Helix” and “S6” predicted in KCNQ1-KCNE3-CaM-PIP2 complex. Only the alpha carbons of the residues on the paths are indicated and coloured by individual amino acids.

After simulations reached convergence (Supplementary Fig. 1 and Supplementary Table 1), we characterized the inactivation paths coupling the VSD at the level of helix S4 to the SF ^42, 43^ via network analysis. Within this framework, the protein was represented as a graph, where nodes are the protein residues and edges are the interactions between residues. A weight was assigned to each edge, which is related to the correlated motions of residues in contact, as computed from MD simulations; this quantity characterizes the efficiency of the motion propagation between the given pair of these residues. Once the network was built, Dijkstra’s algorithm ^44^was used to reveal the communication paths between the VSD and the SF. In the KCNQ1-CaM system, we identified three main inactivation paths: “S1” which passes through S4_n_→S1_n_→P-Helix_n+1_→SF_n/n+1_ (Fig. 1b), “P-Helix” involving S4_n_→S5_n+1_→P-Helix_n+1_→SF_n/n+1_ (Fig. 1c), and “S6” characterized by S4_n_→S5_n+1_→S6_n+1_→SF_n_ (Fig. 1d). The minimal path length d_min_, which is a logarithmic measure of information transfer, is similar among the different paths revealing that their communication efficiency is similar. However, the frequency (f) of each family, computed across all system subunits of all replicas, suggests that the most probable one is “P-Helix” (f=60%), where both helix S5 and the P-Helix act as hubs for the VSD-SF coupling mechanism (Supplementary Table 2). In the KCNQ1-KCNE3-CaM-PIP2 system, the inactivation paths “P-Helix” and “S6” are preserved but become less frequent, while “S1” disappears, suggesting that PIP2 influences inactivation at the level of the VSD-pore domain (PD) interface (Fig. 1e-f, Supplementary Table 2).

The role of each residue in the paths was quantified by computing its betweenness centrality, as reported in the Supplementary Information. This quantity is defined as the number of times a node lies on a minimal length path between the chosen sources and sink regions (Methods and Supplementary Tables 3 and 4). Thus, residues with high betweenness, frequently present in the paths of both systems and acting as key allosteric hubs, were targeted for site-directed mutagenesis to assess their functional roles. Subsequently, their individual contribution on the paths was studied by whole-cell patch-clamp recording. Firstly, we focused on residues of S4 helix, the source in the network, which were mutated to tryptophan to produce marked perturbations ^45^. Then, we selected the most important residues, i.e., exhibiting the greatest values of betweenness, on S1, S5, and P-Helix which were mutated using an alanine-scanning mutagenesis in order to perturb only locally the paths ^46^. Since L273A was non-functional, mutant L273F was used, a known long QTS mutation ^47^. If the mutation interrupts the path, inactivation will be suppressed. Alternatively, if the mutation enhances the coupling of the mutated residue with its neighbors, inactivation will be strengthened. In both cases, the difference from the wild type is expected to be larger, reflecting the prominence of the residue in the path. The impact of each mutation was thus quantified computing the inactivation free energy ΔG_i_, as well as the energetic perturbation of each mutant on the free energy of the inactivation process ΔΔG_i_ as compared to the wild type (see Methods; Tables 1 and 2).

**Table 1.**
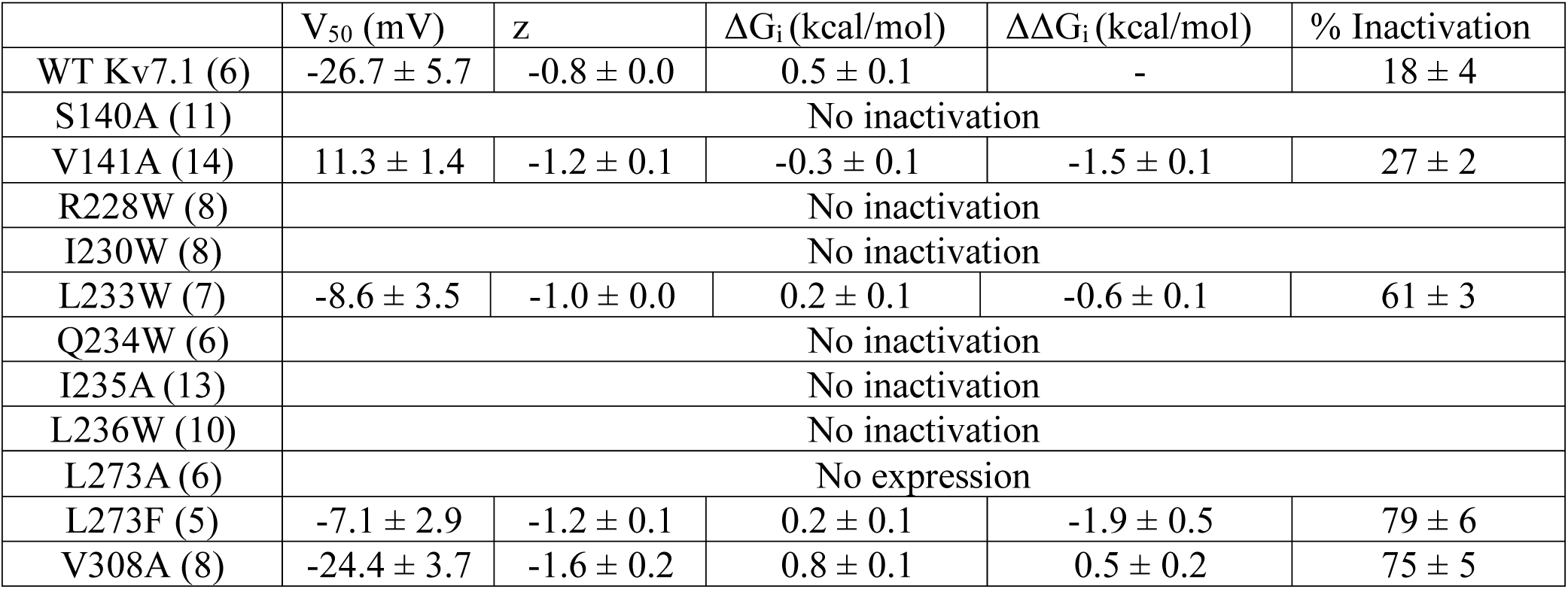
Fitted values of steady-state voltage dependence of inactivation and their deduced free energy of WT Kv7.1 and mutants along the VSD-SF inactivation paths. Values in parentheses correspond to the number of independent recorded cells. The percentage of inactivation was measured at +60 mV. The activation values of these mutations are reported in supplementary Table 8.

| | $V_{50}$ (mV) | $z$ | $\Delta G_i$ (kcal/mol) | $\Delta\Delta G_i$ (kcal/mol) | % Inactivation |
| --- | --- | --- | --- | --- | --- |
| WT Kv7.1 (6) | $-26.7 \pm 5.7$ | $-0.8 \pm 0.0$ | $0.5 \pm 0.1$ | - | $18 \pm 4$ |
| S140A (11) | No inactivation |  |  |  |  |
| V141A (14) | $11.3 \pm 1.4$ | $-1.2 \pm 0.1$ | $-0.3 \pm 0.1$ | $-1.5 \pm 0.1$ | $27 \pm 2$ |
| R228W (8) | No inactivation |  |  |  |  |
| I230W (8) | No inactivation |  |  |  |  |
| L233W (7) | $-8.6 \pm 3.5$ | $-1.0 \pm 0.0$ | $0.2 \pm 0.1$ | $-0.6 \pm 0.1$ | $61 \pm 3$ |
| Q234W (6) | No inactivation |  |  |  |  |
| I235A (13) | No inactivation |  |  |  |  |
| L236W (10) | No inactivation |  |  |  |  |
| L273A (6) | No expression |  |  |  |  |
| L273F (5) | $-7.1 \pm 2.9$ | $-1.2 \pm 0.1$ | $0.2 \pm 0.1$ | $-1.9 \pm 0.5$ | $79 \pm 6$ |
| V308A (8) | $-24.4 \pm 3.7$ | $-1.6 \pm 0.2$ | $0.8 \pm 0.1$ | $0.5 \pm 0.2$ | $75 \pm 5$ |

**Table 2.** Fitted values of steady-state voltage dependence of inactivation and their deduced free energy of WT Kv7.1 and mutants along the CaM-SF allosteric paths. Values in parentheses correspond to the number of independent recorded cells. The percentage of inactivation was measured at +60 mV. nd, not determined. The activation values of these mutations are reported in supplementary Table 8.

| System | $V_{50}$ (mV) | $z$ | $\Delta G_i$ (kcal/mol) | $\Delta\Delta G_i$ (kcal/mol) | % Inactivation |
| --- | --- | --- | --- | --- | --- |
| WT Kv7.1 (6) | $-26.7 \pm 5.7$ | $-0.8 \pm 0.0$ | $0.5 \pm 0.1$ | - | $18 \pm 4$ |
| C180A (9) | nd | nd | nd | nd | $2.5 \pm 1.0$ |
| R181A (11) | No inactivation |  |  |  |  |
| S182A (4) | $-41.9 \pm 2.9$ | $-1.4 \pm 0.3$ | $1.5 \pm 0.3$ | $10.6 \pm 3.8$ | $44 \pm 2$ |
| Y184F (11) | nd | nd | nd | nd | $35 \pm 4$ |
| L239W (5) | No inactivation |  |  |  |  |
| Q244W (9) | nd | nd | nd | nd | $39.3 \pm 3.1$ |
| I274A (6) | $8.9 \pm 3.4$ | $-1.0 \pm 0.1$ | $-0.2 \pm 0.1$ | $-0.9 \pm 0.1$ | $9 \pm 1$ |
| G306A (6) | No inactivation |  |  |  |  |
| V310G (9) | $-24.2 \pm 3.8$ | $-1.3 \pm 0.3$ | $0.8 \pm 0.3$ | $0.4 \pm 0.3$ | $87 \pm 3$ |
| T312S (8) | $-31.7 \pm 3.7$ | $-1.0 \pm 0.1$ | $0.8 \pm 0.1$ | $0.8 \pm 0.4$ | $28 \pm 5$ |
| S338F (10) | $12.8 \pm 10.7$ | $-1.4 \pm 0.2$ | $-0.4 \pm 0.2$ | $-0.9 \pm 0.3$ | $59 \pm 2$ |
| F351A (8) | No inactivation |  |  |  |  |
| K354W (7) | No inactivation |  |  |  |  |
| Q357W (13) | No inactivation |  |  |  |  |
| R366W (11) | $-32.2 \pm 16.4$ | $-1.1 \pm 0.4$ | $0.8 \pm 0.2$ | $1.4 \pm 0.2$ | $77 \pm 4$ |
| WT Kv7.1+WT CaM (10) | nd | nd | nd | nd | $13.3 \pm 2.2$ |
| WT Kv7.1+ N98R CaM (8) | No inactivation |  |  |  |  |
| WT Kv7.1+Y100A CaM (7) | No inactivation |  |  |  |  |
| WT Kv7.1+Y100D CaM (6) | No inactivation |  |  |  |  |

All tested residues affected inactivation either by improving or impairing the inactivation transition. Two main gating phenotypes were identified. (i) non-inactivating mutants such as S140A in S1, R228W, I230W, Q234W, I235W, L236W in S4 (paths “S1”, “S6”, “P-helix”), which do not inactivate. These mutational perturbations interrupt the inactivation paths, such that the inactivated state is destabilized (Fig. 2a,; Table 1). For example, the inactivation path jumps from R231 in S4 to S140 in S1 (route “S1”) of the same subunit. The R231-S140 interaction corresponds to an H-bond between the side chain of S140 and the backbone carbonyl of R231 that is disrupted by the S140A mutation. As a result, the “S1” inactivation path is interrupted, and the phenotype of the mutant is non-inactivating (Fig. 2). Another example shows that for residue I235, the inactivation path route “P-Helix” jumps from I235 on S4 to L273 on helix S5 of the neighboring subunit (n+1). In this case, the I235-L273 hydrophobic interaction could possibly be disrupted when I235 is mutated to the very bulky tryptophan residue. This leads to an interruption of this path with a resulting non-inactivating phenotype. Notably, the “P-Helix” inactivation path is the most frequent route, so the mutational impact is expected to be strong. (ii) inactivating mutants with larger inactivation than WT such as V141A in paths “S1”, L233W in S4, L273F in S5 and V308A in path “P-Helix” and where the mutational perturbations stabilize the inactivated state (Fig. 2b-d; Table 1). For example, the inactivation path route “P-Helix” jumps from L236 on S4 to L273 on S5 and converges to the SF via the P-Helix V308. L273 is at atomic proximity with crucial residues of the P-Helix L303, G306, V307 and V310, which stabilize the conducting conformation of the SF. Mutating L273 with the aromatic phenylalanine may destabilize the SF and triggers inactivation. Another example concerns residue V308 at the end of the P-Helix at the point where the route “P-Helix” converges to the SF, at the level of Y315. V308 is at atomic proximity with critical residues such as W304 or Y315 that stabilize the SF and replacing the bulky valine with the smaller alanine may destabilize the conducting filter conformation and produce inactivation. Taken together, these results confirm the theoretical predictions on the inactivation paths, demonstrating the role of helices S4, S5, and P-Helix in coupling the VSD to the SF.

**Figure 2:**
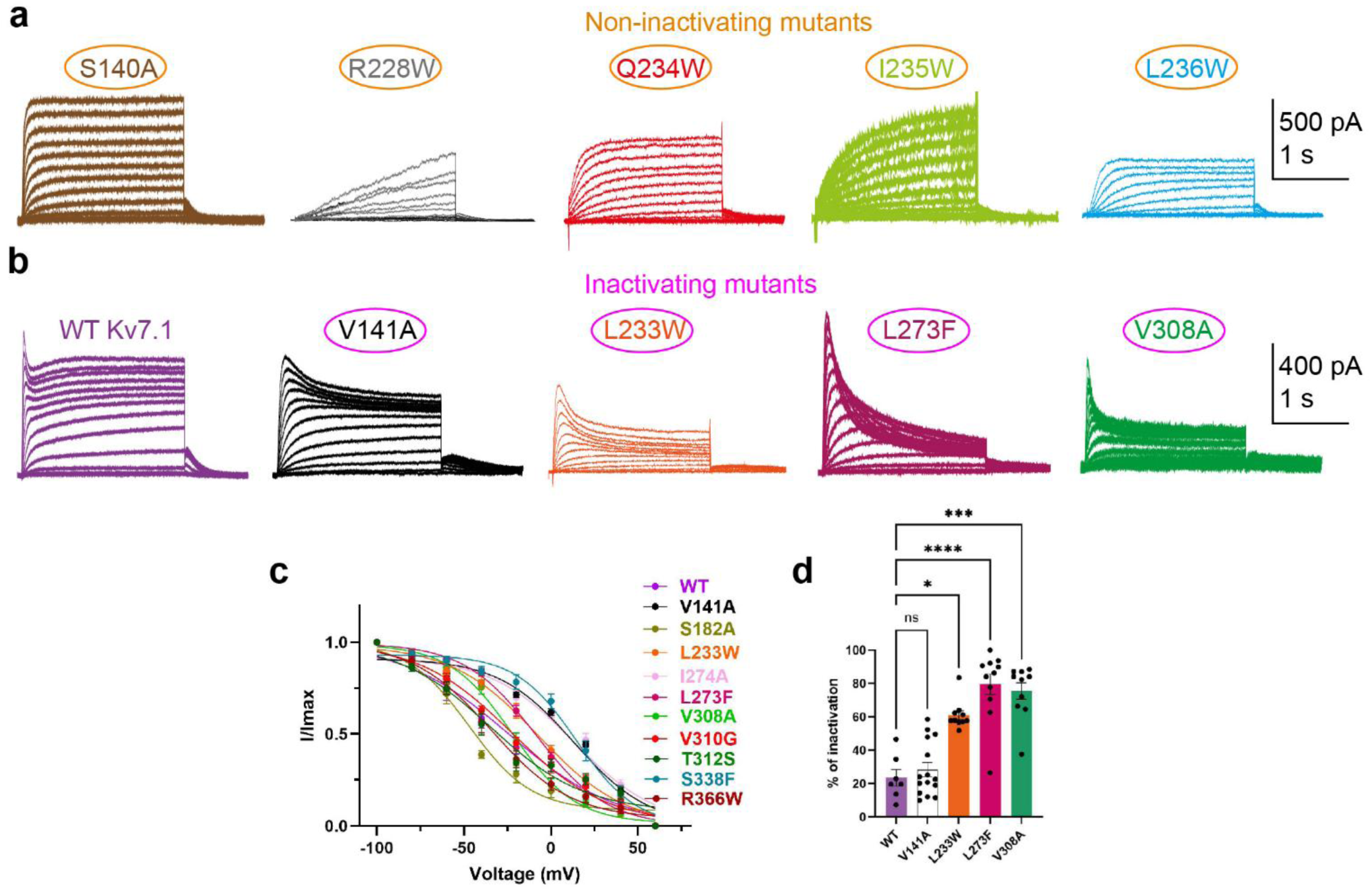
Properties of the Kv7.1 mutants along the VSD-SF inactivation paths. **a.** representative traces of the non-inactivating mutants, S140A, R228W, Q234W, I235W and L236W. Recordings were performed in 1.8 mM external Ca2+ solutions and in nominally 0 µM free Ca2+ internal pipette solution. From a -90 mV holding potential, a series of depolarizing voltage steps from -60 to +60 mV, in 10 mV increments were applied for 3 s, followed by a repolarization to -60 mV for 1.5 s. **b.** representative traces of WT Kv7.1 and the inactivating mutants, V141A, L233W, L273F and V308A. **c.** Boltzmann fitting of normalized steady-state inactivation curves as a function of the inactivating pre-pulse voltage of WT Kv7.1 and V141A, S182A, L233W, I274A, L273F, V308A, V310G, T312S, S338F, R366W. The fitted values (V_50_ and z) are displayed in Table 1; data are expressed as mean ± SEM. From a -90 mV holding potential, a series of pre-pulse depolarizing voltage steps from -100 to +60 mV, in 20 mV increments were applied for 2 s to allow the channels to open and reach a steady inactivated state. A test pulse to +60 mV was applied for 300 ms to measure the remaining available current. This test pulse activates the channels that were not inactivated during the pre-pulse. **d**. Inactivation of WT Kv7.1 (23.5 %, n=7) and the mutants V141A (28.4 %, n=15), L233W (61%, n=10), L273F (79.5%, n=11) and V308A (75.4%, n=10). Inactivation was quantified from the ratio current of the lowest current amplitude after the decay of the transient component (dip) and the peak amplitude at +60 mV and expressed as percentage. One-way ANOVA, Kruskal-Wallis’s test; *P<0.05, ***P=0.0005 and ****P<0.0001.

### CaM-SF regulation paths

The communication mechanism between CaM and the SF was subsequently investigated to understand how Ca^2+^-CaM regulates the inactivation. In the KCNQ1-CaM system, six families of regulation paths were identified (Fig. 3a-f): CaM_n_→S1_n_→S5_n+1_→SF_n+1_ and CaM_n_→S1_n_→H45_n_→S5_n+1_→P-Helix_n+1_→SF_n_ defining the so-called “S0-S1” and “S0-H45” paths, where the information is transferred from CaM to the channel via the bend between helices S0 and S1; CaM_n_→S2_n_→S4_n_→S5_n+1_→SF_n_ and CaM_n_→S3_n_→S4_n_→P-Helix_n+1_→SF_n+1_ where the paths reach the channel traversing the S2-S3 loop (“S2” and “S3” paths); finally, CaM_n_→HB_n_→S6_n_→S6_n+1_→SF_n_ and CaM_n_→HA_n_→S6_n_→S5_n+1_→S6_n+1_→P-Helix_n+1_→SF_n_ moving from CaM to the channel through helices HA or HB (“HA” and “HB” paths) (Fig. 3a-f). The frequency of such paths reveals that those traversing helices S3 and HB are the most frequent (Supplementary Table 2 and Supplementary Tables 5-7), thus being crucial for the CaM-SF coupling mechanism. Interestingly, paths “S0-S1”, “S0-H45”, and “S3” all converge on the P-Helix, joining inactivation paths “S1” and “P-Helix”. In the KCNQ1-KCNE3-CaM-PIP2 system (Fig. 3g), only path “HA” was found between CaM and SF, due to the structural rearrangements occurring at the CaM in the presence of PIP2 that disconnect CaM from the VSD ^40^. Overall, these results indicate that the CaM-SF allosteric paths share significant overlaps with the inactivation paths, converging mainly via the routes involving S5 helix and P-Helix.

**Figure 3:**
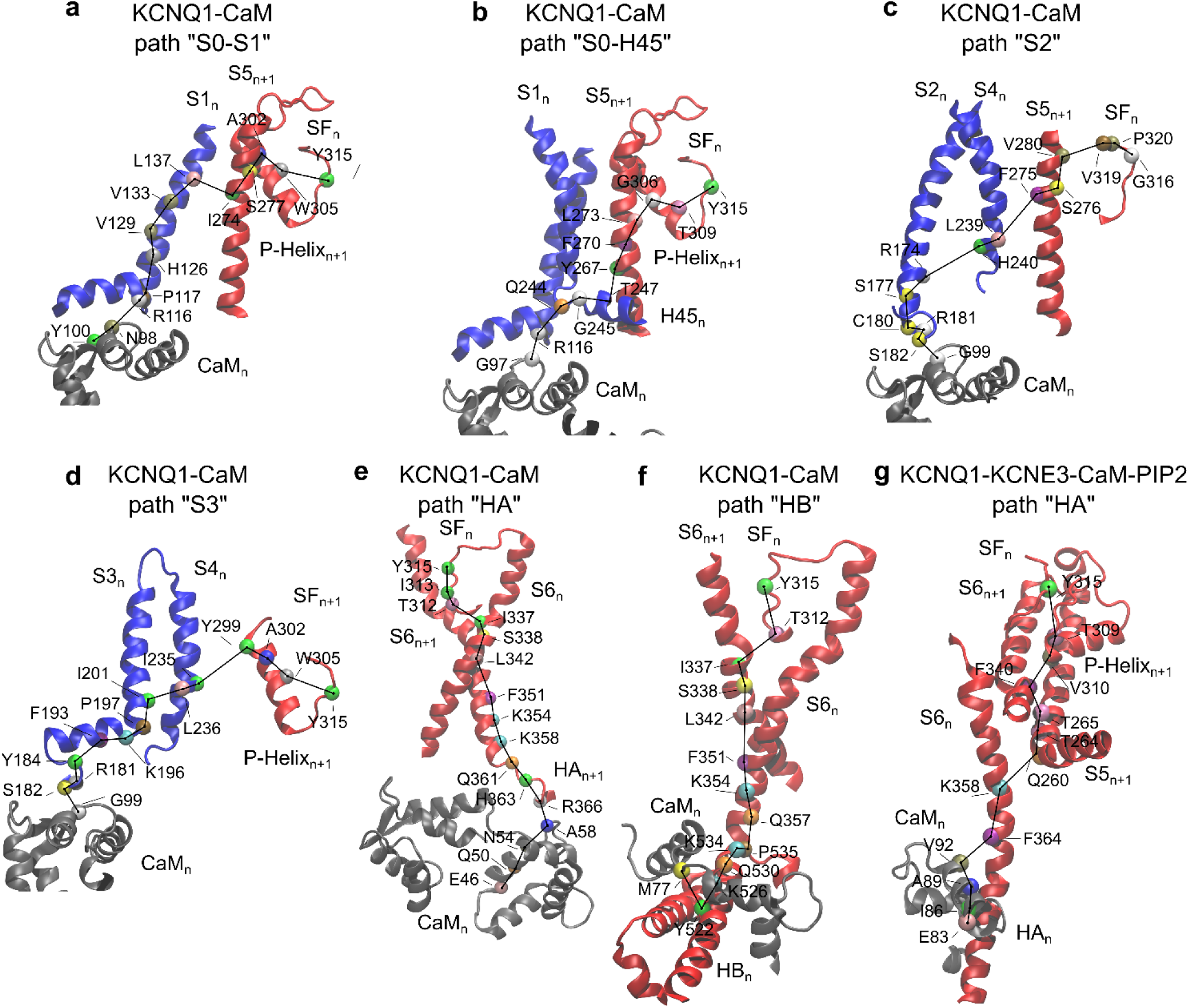
Molecular basis for the CaM-SF regulation paths. Panels **a-f** show the CaM-SF paths “S0-S1”, “S0-H45”, “S2”, “S3”, “HA” and “HB” predicted in the KCNQ1-CaM system. **g** CaM-SF “HA” path identified in the KCNQ1-KCNE3-CaM-PIP2 complex. Only the alpha carbons of the residues on the paths are indicated and coloured by individual amino acids.

As for the inactivation paths, we mutated residues lying on the CaM-SF paths to confirm such theoretical predictions (Fig. 4, Supplementary Tables 6-9). Also in this case, two outcomes are expected: since the function of these paths is to repress the inactivation, a mutation interrupting the path will lead to worsened inactivation, while a mutation increasing the efficiency of signal propagation along the path will produce a non-inactivating phenotype. Thus, the experimental validation along the CaM-SF paths revealed two main gating behaviors: (i) non-inactivating mutants such as R181A on S2-S3 loop, L239W on S4 (CaM-SF paths “S2” and “S3”), G306A, F351A, K354W, and Q357W on S6 (CaM-SF path “HB”) (Table 2 and Fig. 4; (ii) inactivating mutants with larger inactivation than WT such as S182A and Y184F on S2-S3 loop, Q244W on S4-S5 loop (CaM-SF path “S0-H45”), I274A on S5 (CaM-SF path “S0-S1” and “S0-H45”), V310G on P-Helix (CaM-SF paths “S0-H45” and “HA”), T312S on SF, the two LQTS-linked mutants S338F on S6 and R366W on HA helix (CaM-SF paths “HA” and “HB”) (Table 2 and Fig. 4). C180A, instead, shows significantly lower inactivation than WT.

**Figure 4:**
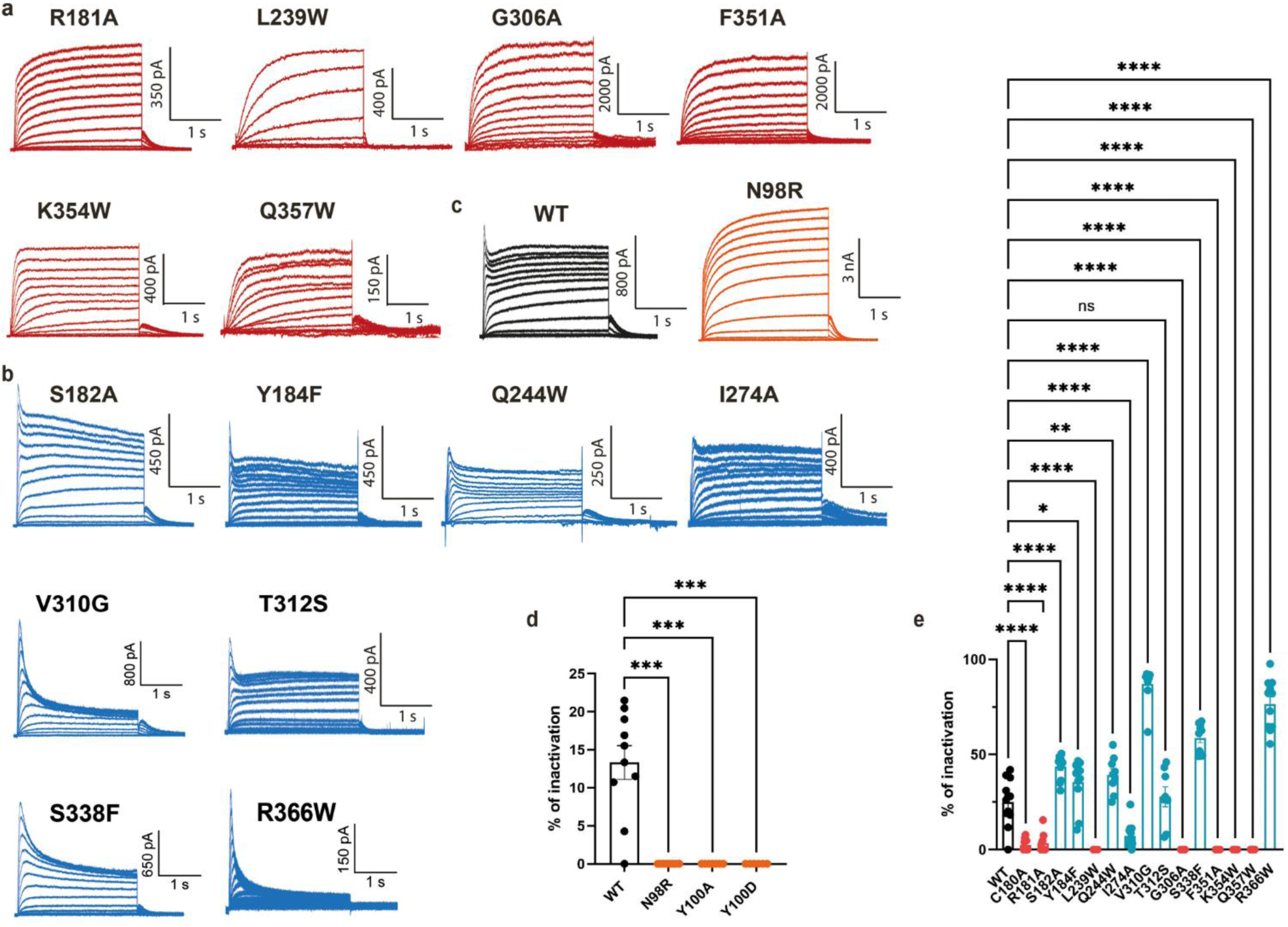
Properties of the Kv7.1 mutants along the CaM-SF regulation paths. **a.** Representative traces of the non-inactivating Kv7.1 mutants (red traces) R181A, L239W, G306A, F351A, K354W and Q357W. The current voltage protocol was the same as that described in Figure 2. **b.** Representative traces of the inactivating Kv7.1 mutants (blue traces) S182, Y184F, Q244W, I274A, V310G, T312S, S338W and R366W. **c.** Representative traces of WT Kv7.1 (black trace) and of the CaM mutant N98R (orange trace) co-expressed with WT Kv7.1. **d.** Inactivation of WT (13.3 %, n=10) and CaM mutants N98R (0%, n=8), Y100A (0%, n=7) and Y100D (0 %, n=6) co-expressed with WT Kv7.1. One-way ANOVA, Kruskal-Wallis test; ***P=0.0002 for N98R, ***P=0.0003 for Y100A and ***P=0.0006 for Y100D; recording was performed in external Ca^2+^ containing solutions and in nominally 0 µM free Ca^2+^ internal solution. **e.** Inactivation of WT Kv7.1 (25.1%, n=12) and its mutants C180A (2.5%, n=9), R181A (3.5%, n=9), S182A (43.5%. n=11), Y184F (35.5%, n=11), L239W (0%, n=5), Q244W (39.3%, n=9), I274A (7.2%, n=12), G306A (0%, n=6), V310G (87.0%, n=9), T312S (27.8%, n=8), S338F(58.7%, n=10), F351A (0%, n=8), K354W (0%, n=7), Q357W (0%, n=13) and R366W (76.6%, n=11). One-way ANOVA, Dunnett’s multiple comparisons test, ****P<0.0001for WT vs C180A, R181A, S182A, L239W, I274A, G306A, V310G, S338F, F351A, K354W, Q357W and R366W; **P=0.0030 for WT vsQ244W, *P=0.0460 for WT vs Y184F and ns (not significantly different) for WT vs T312S.

Similarly, mutants on CaM with the highest values of betweenness were studied by co-expressing the channel with CaM, with the objective to interrupt the CaM-SF paths at the level of their starting points: N98R, Y100A, and Y100D (Fig. 4, Table and Supplementary Table 7). All these mutants display a significantly weaker inactivation than WT CaM. Taken together, these results demonstrate that mutations on the CaM-SF paths and not lying on the inactivation paths, can significantly influence inactivation with either positive or negative effects as compared to WT, thus demonstrating that the communication between CaM and SF plays a major role in regulating inactivation.

### Comparison between theoretical predictions and experimental analyses

To quantitatively compare the theoretical predictions and the experimental results, we plotted the betweenness centrality of each mutated residue computed by network analysis against the absolute value of ΔΔG_i_; this scatter plot reveals correlations between the importance of a residue along a path and its effect on a given gating process ^41^. The betweenness centrality of residues involved in the inactivation paths have correlation coefficients R=0.57 and R=0.62 in KCNQ1-CaM and KCNQ1-KCNE3-CaM-PIP2 structures, respectively (Fig. 5). These correlation coefficients are comparable to those obtained for hERG inactivation in our previous work ^41^. The correlation is generally poorer for inactivation than for activation possibly due to the weaker steepness of the voltage dependence of inactivation (smaller values and variations of the gating charge z_i_) and larger relative errors in ΔΔG_i_. Interestingly, when comparing the betweenness centrality of the residues involved in the CaM-SF paths with ΔΔG_i_, a strong correlation was observed in the KCNQ1-CaM system (R=0.86). On the other hand, in the KCNQ1-KCNE3-CaM-PIP2 system, no correlation was found (R=0.01, Fig. 5f), since CaM communicates with the channel only through the “HA” path. As a result, all residues on the intracellular side show very low betweenness values, as they are not part of other paths to reach the channel. Overall, these data demonstrate a fair agreement between computational and experimental results, reinforcing the notion that the identified residues are part of the inactivation and of the CaM-SF paths in Kv7.1 channels.

**Figure 5:**
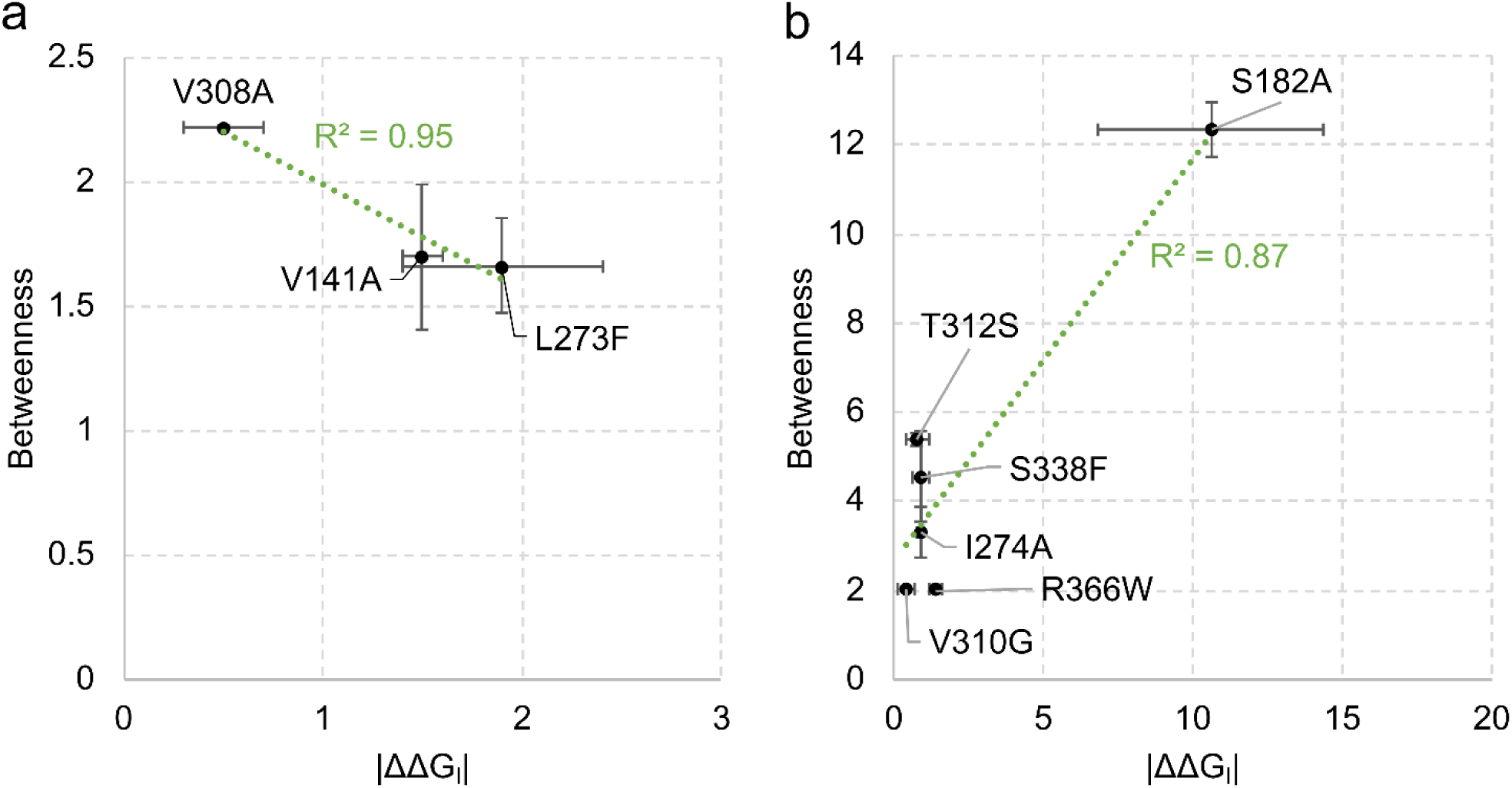
Scatter plots comparing the betweenness centrality of WT residues along the inactivation and the CaM-SF paths against free-energy perturbation of inactivation (ΔΔG_i_). **a** Betweenness centrality of residues along the inactivation paths vs ΔΔG_i_. **b** Betweenness centrality of residues along the CaM-SF paths vs ΔΔG_i_. Green lines are linear fits to the data computed by the regression analysis and “R” is the corresponding correlation coefficient. Data are presented as mean values ± standard error.

## Discussion

In comparison to most voltage-gated K^+^ channels, Kv7.1 features a unique gating machinery, in which its interaction with CaM plays a pivotal role in both activation and inactivation ^35, 39, 40, 48^. The goals of this study were to characterize the allosteric paths leading to Kv7.1 inactivation and to identify the allosteric mechanisms coupling the Ca^2+^-CaM signal from its intracellular site to the SF, which prevents channel inactivation.

Just as in an urban setting there are many alternative routes to go from point A to point B, we found different path families connecting our initial (source) and target points (sink). For instance, we identified three families of inactivation paths. Notably, despite differences in their initial source, they tend to converge on the S5 segment and P-Helix before finally reaching the SF, as confirmed by our experiments on the non-inactivating mutants R228W, I230W, Q234W, I235W, and L236W, whose mutations modified the inactivation process. These paths resemble the inactivation pathways previously predicted in the domain-swapped Kv1.2 channels ^49^. Interestingly, those passing through helix S1 – termed "S1" inactivation paths – resemble the noncanonical paths found in a different class of potassium channels, the non-domain-swapped hERG channels, thus suggesting an evolutionary remnant underlying this path conservation ^41^. However, the conformational changes that trigger Kv7.1 inactivation remain elusive for at least two reasons: first, an inactivated structure of Kv7.1 is not yet available, and second, Kv7.1 inactivation is different from both the N-type and C-type mechanisms. Structural studies of an inactivated KcsA channel showed a constriction of the SF ^7, 8^. However, the relevance of this constricted conformation to C-type inactivation in Kv channels has been debated ^9, 10, 11, 12^. A recent structure of the Kv1.2 channel revealed that C-type inactivation involves dilation of the SF that disrupts the outer two K^+^ binding sites S1 and S2 ^50^. The changes at the SF propagate to the extracellular pore mouth and the turret regions of the channel ^50^. Interestingly, structural and simulation studies showed that the K2P2.1 channel (TREK-1) undergoes asymmetric order-disorder transitions, encompassing pinching and dilation of the SF, thereby disrupting the outer S1 and S2 K^+^ binding sites ^51^. Clear differences exist between the C-type inactivation of *Shaker*-like Kv channels and Kv7.1 inactivation, such as the role of external K^+^, the coordination of Ca^2+^ in the turret region, and the striking regulation by PIP2 and Ca^2+^-CaM. However, other important features are shared and converge at the SF. Notably, the conserved aspartate at the filter entrance and tryptophan at the P-Helix, along with their H-bonding network behind the filter, suggest that SF inactivation is an ancient process arising from the shared voltage-gated K^+^ channel pore architecture ^35, 52, 53^. Future structural studies should provide a comprehensive picture of the SF conformation underlying the unique Kv7.1 channel inactivation.

Here, we demonstrated the presence of six families of paths coupling CaM to the SF. The contact points between CaM and the channel were found at the level of helices S0, S1, S2, S3, H45, HA, and HB. Interestingly, the S2-S3 linker and HA-HB helices have been previously identified to interact with CaM in modulating Kv7.1 voltage-dependent gating mechanisms, thus revealing that they are crucial for regulating both activation and inactivation of Kv7.1 channels ^35, 39, 48^. In agreement with our theoretical predictions, mutations R181A, L239W, G306A, F351A, K354W, and Q357W on the Kv7.1 channel side of the Kv7.1/CaM interface yielded a non-inactivating phenotype, while mutations S182A, Y184F, I274A, V310G, T312S, S338F and R366W, produced a strong inactivating effect. Finally, the CaM mutations N98R, Y100A, and Y100D caused weaker inactivation than WT CaM. There is a structural basis for the intimate interaction between the CaM C-lobe residues N98 and Y100 and those of the S2-S3 channel linker, R181 and S182 ^39^. Indeed, in the "attached CaM configuration", all these residues are in atomic proximity. N98 can engage in H-bonding with the guanidinium of R181; similarly, Y100 can interact via H-bonding with S182 ^39^. This tight and intricate network of interactions confirms that the interface between the CaM C-lobe and the VSD S2-S3 linker plays a crucial role in the Kv7.1 gating process and is involved in the allosteric network used by the CaM signal to preserve the selectivity filter in a conducting conformation and prevent inactivation.

Altogether, our combined experimental and simulation findings reveal that the mechanism of Kv7.1 voltage-dependent inactivation can be understood in terms of VSD-SF communication, which enables the allosteric coupling between the voltage sensor and the selectivity filter. The inactivation paths involve the S4/S1, S4/S5, and S5/S6 subunit interfaces, all converging to the P-Helix and the SF as an endpoint for inactivation.

Moreover, our study also identifies novel CaM-SF allosteric paths via the S5 segment and the P-helix that underlie the mechanism by which the Ca^2+^-CaM signal emerges from its intracellular boundary to converge to the SF and prevent Kv7.1 inactivation. Importantly, these new allosteric communication paths may also contribute to understand the regulation of other ion channels, whose gating is controlled by CaM.

## Materials and Methods

### Constructs

Human Kv7.1 and rat CaM were cloned into the pcDNA3 vector to allow eukaryotic expression. The mutations in Kv7.1 and in CaM were introduced using the PCR-based Quikchange site-directed mutagenesis (Stratagene) and were verified by full sequencing of the entire plasmid vector.

### Cell Culture and Transfection

Chinese hamster ovary CHO cells were grown in Dulbecco’s modified Eagle’s medium supplemented with 2 mM glutamine, 10% fetal calf serum, and antibiotics. In brief, 40,000 cells seeded on poly-D-lysine-coated glass coverslips (13 mm in diameter) in a 24-multiwell plate were transfected with pcDNA3 EGFP (0.18 µg) as a fluorescent marker for transfection and with 0.5 µg WT Kv7.1 and its mutants or WT CaM and its mutants. Transfection was performed using 1.75 µl of TransIT-LT1 Transfection Reagent (Zotal) according to the manufacturer’s protocol. For electrophysiology, transfected cells were visualized approximately 40h after transfection, using a fluorescence microscope (Nikon Eclipse Ts2) with green-fluorescent filter.

### Preparation of computational systems

The human KCNQ1-CaM system was produced from the experimentally solved structure (PDB ID: 6UZZ) ^39, 40^ and the missing loops were modelled with MODELLER ^54^. Since the gap 397-505 of a cytosolic domain was too long to be predicted by homology modelling, residues I396 and L506 were capped with a N-methyl and an acetyl group, respectively, to avoid spurious electrostatic interactions. The APBS software ^55^ was used to predict the protonation states of each residue. All aspartates and glutamates were ionized. Histidines 105, 126, 258, 509, 510, and 549 were predicted to be in the δ protonation state while histidines 108, 240, and 363 were assigned to the ε state. Using the CHARMM Membrane Builder ^56, 57^, the ion channel was embedded into a 520 POPC lipid bilayer and solvated with 79,027 TIP3P water molecules and 0.15 M of NaCl, totaling 339,411 atoms; the box size was ca. 164×161×164 Å.

The human KCNQ1-KCNE3-CaM complex with PIP2 was produced from the experimental structure (PDB ID: 6V01) ^40^ and the missing loops were modelled with MODELLER ^54^. Residues P387 and L506 were capped with a N-methyl and an acetyl group due to the presence of a missing loop too long to be modelled with state-of-art tools between residues 388-505. The APBS software ^55^ was used to predict the protonation states of each residue. All lysines and arginines were protonated while aspartates and glutamates were ionized except for D57 and E295 which were protonated. Histidines 105, 108, 126, 240, 363, 509, 510 were predicted to be in the δ state; H44 was assigned to δ state in two subunits and to ε state to the others; H258 was assigned to P state. The CHARMM Membrane Builder ^56, 57^ was used to embed the ion channel in a 534 POPC lipid bilayer with 76,218 TIP3P water molecules and 0.15 M of NaCl, totalling 335,190 atoms; the box size was ca. 155×157×168 Å.

### MD simulations

Three independent replicas per system have been run for more than 1 μs with NAMD 2.14 ^58^ using the ff19SB force field for the protein ^59^, the Lipid21 force field for the lipids ^60^ and the TIP3P water model ^61^. The parameters from Künze lab have been used for the PIP2 ^62, 63^, while those for calcium were taken from Bradbrook et al ^64^. In all simulations, pressure was kept at 1.01325 bar by the Nosé-Hoover Langevin piston method ^65^ and the temperature was maintained at 303.15 K by a Langevin thermostat with damping coefficient of 1 ps^-1^. Long-range electrostatic interactions were evaluated with the smooth PME algorithm ^66^ with a grid space of 1 Å. For short-range non-bonded interactions, a cut-off of 12 Å with a switching function at 10.0 Å was used. The integration time step was 2 fs. The stability of the systems during the dynamics was monitored by computing the Root Mean Square Deviation (RMSD) of the backbone atoms of the protein from the initial structure. The trajectories were visually inspected using VMD 1.9.3 software ^67^.

### Network analysis

The communication paths between the voltage sensor domain at the level of helix S4 and CaM to the SF were studied by employing a network analysis with a graph representation of the protein ^68^. The weight assigned to each graph’s edge was:

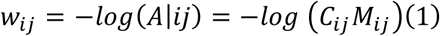

being *C*_*ij*_ the semi-binary contact map and *M*_*ij*_ the matrix that quantifies the motion correlation of residues.

*C*_*ij*_ was computed with a truncated Gaussian kernel:

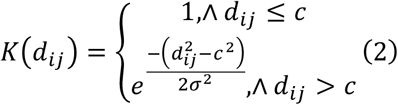

where *d*_*ij*_ is the distance between the *C*_*α*_ of the *i* and *j* residues and *c* is the cut-off distance set to 7.0 Å. The width *σ* of the Gaussian kernel was chosen so as to attain a negligibly small value of the kernel at *d*_*ij*_ = 10Å. Specifically, we imposed *K*(*d*|*cut*) = 10^−5^ attaining *σ* = 1.48. The final contact map was computed by averaging the value of the kernel over all the frames of the trajectory:

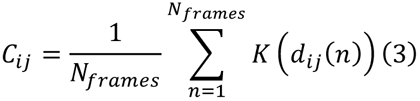

*M*_*ij*_ was computed as:

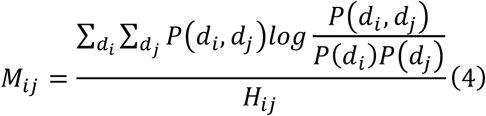

where *d*_*i*_ and *d*_*j*_ are the displacement of the center of mass of the side chain of the *i* and *j* residues with respect to its average position and *H*_*ij*_ is the Shannon entropy ^69^ of those variables.

Dijkstra’s algorithm ^44^ was used to compute the minimal paths between helix S4 and CaM of the *n* subunit (source) to the SF of the same (*n*, intra-subunit paths) or the neighboring (*n* + 1, inter-subunit paths) subunit (sink). All the residues of helix S4, CaM and SF were considered in the network. The *d*_*min*_ value corresponds to the mean over the four subunits for each replica ± the standard deviation of the lowest value computed from Eq. 1 between each residue of the source and each residue of the sink regions. The frequency of each family of path was computed as: 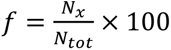 where *N*_x_ is the number of paths passing through helix *x* and *N*_*tot*_ is the total number of paths coupling the source to the sink region. Betweenness centrality is defined as a measure of how often a node lies on a path of minimal length connecting the chosen source and sink regions in a network. The betweenness centrality of each residue was computed with Brandes’s algorithm ^70^ as implemented in the NetworkX 3.0 library ^71^.

### Electrophysiology

Recordings were performed using the whole cell configuration of the patch clamp technique. Signals were amplified using an Multiclamp 700B patch-clamp amplifier (Axon Instruments), sampled at 10 kHz and filtered at 4 kHz via a four pole Bessel low pass filter. Data were acquired using pClamp 10. 7 software in conjunction with a DigiData 1440A interface. The patch pipettes were pulled from borosilicate glass (Harvard Apparatus) with a resistance of 3–5 megaohms. The intracellular pipette solution contained 130 mM KCl, 5 mM K_2_-ATP, 10 mM HEPES, pH 7.3 (adjusted with KOH) and 5 mM K4BAPTA (no added CaCl_2_ for nominally 0 free Ca^2+^ concentration as calculated by MAXCHELATOR (WEBMAXC STANDARD) software; https://somapp.ucdmc.ucdavis.edu/pharmacology/bers/maxchelator/webmaxc/webmaxcS.htm), with sucrose added to adjust osmolarity to 290 mosmol. The external solution was 140 mM NaCl, 4 mM KCl, 1.8 mM CaCl_2_, 1.2 mM MgCl_2_, 11 mM glucose, 5.5 mM HEPES, adjusted with NaOH to pH 7.3 (310 mOsM). Series resistances ranged between 8 and 12 MΩ and were compensated for V-clamp recording using the adjust bottom Rs of the Multiclamp 700B amplifier (providing ≈85–90% compensation). Electrophysiological data analysis was performed using the Clampfit program (pClamp 10.7; Axon Instruments), Microsoft Excel (Microsoft,Redmond, WA), and Prism 9.0 (GraphPad Software, Inc., San Diego, CA). Leak subtraction was performed off-line, using the Clampfit program of the pClamp 10.7 software. Chord conductance (G) was calculated by using the following equation: G=I/(V-Vrev), where I corresponds to the current amplitude measured at the end of the pulse, and V_rev_ is the calculated reversal potential assumed to be -90 mV in CHO cells. G was estimated at various test voltages (V) and then normalized to a maximal conductance value, Gmax. Activation curves were fitted by one Boltzmann distribution: G/Gmax=1/{1+exp[(V_50_-V)/s]}, where V_50_ is the voltage at which the current is half-activated and s is the slope factor. Inactivation was quantified from the ratio current of the lowest current amplitude after the decay of the transient component (dip) and the peak amplitude at +60 mV and expressed as percentage. Steady-state inactivation curves were fitted by one Boltzmann distribution: I/Imax=1/{1+exp[(V_50_-V)/s]}, where V_50_ is the voltage at which the current is half-activated and s is the slope factor. For the thermodynamic analysis, the difference in Gibbs free energy between closed and open states at 0 mV (ΔG_0_) was calculated according to ΔG_0_ = 0.2389 zFV_50_, where z = RT/Fxs. The difference in free energy for inactivation was calculated according to ΔG_i_ = 0.2389 zFV_50_.The change in free energy difference for activation gating between WT Kv7.1 and the mutant was computed by ΔΔG_0_ = ΔG_0_^mut^-ΔG_0_^wt^. To take into account the differences in side chain volume |Δvol| produced by the mutations, ΔΔG_0_^c^ values were corrected according to the relative change in side chain volume ΔΔG_0_^c^ = ΔΔG_0_ [|Δvol^av^/Δvol|], with an average change vol^av^ = 68 Å^3^, as previously described ^46, 70, 72^. The change in free energy difference for inactivation gating between WT Kv7.1 and the mutant was computed by ΔΔG_i_ = ΔG_i_^mut^-ΔG_i_^wt^. All data were expressed as mean ± S.E.M. Statistically significant differences were assessed by paired, unpaired two-tailed Student’s t test or one-way ANOVA, as indicated in figure legends.

## Acknowledgments

F.C., C.G., and A.G. acknowledge PRACE for awarding us to access to the Leonardo Supercomputer at CINECA, Italy. F.C. and A.G. acknowledge financial support by the Italian Ministry of Education, University, and Research (MIUR) through the “Framework per l’Attrazione e il Rafforzamento delle Eccellenze per la Ricerca in Italia (FARE)” scheme, grant SERENA n. R18XYKRW7J. C.G. and A.G. acknowledge financial support under the National Recovery and Resilience Plan (NRRP), Mission 4, Component 2, Investment 1.1, Call for tender No. 1409 published on 14.9.2022 by the Italian Ministry of University and Research (MUR), funded by the European Union – NextGenerationEU– Project Title “The virtual EV (v-EV): A digital twin of extracellular vesicles for health and food” – CUP B53D23027530001 – Grant Assignment Decree No. 1389 adopted on 01/09/2023 by the Italian Ministry of Ministry of University and Research (MUR). This work was also supported by the Israel Science Foundation grants 3129/21 and 1653/21 and 1806/25 to B.A. and Y.H., respectively, and by the binational Israel-USA Science Foundation 2019159 to B.A.

## Author Contributions

Conceptualization: C.G., A.G., Y.H., B.A. Investigation: N.D., F.C., C.G., M.L., A.B. Supervision: A.G., Y.H., B.A. Writing: F.C., A.G., Y.H., B.A.

## Materials & Correspondence

Yoni Haitin; Alberto Giacomello; Bernard Attali

