## Supplementary information for "Ca^2+^-calmodulin regulates Kv7.1 channel gating by allosterically interfering with its inactivation path"

**Supplementary Figures**

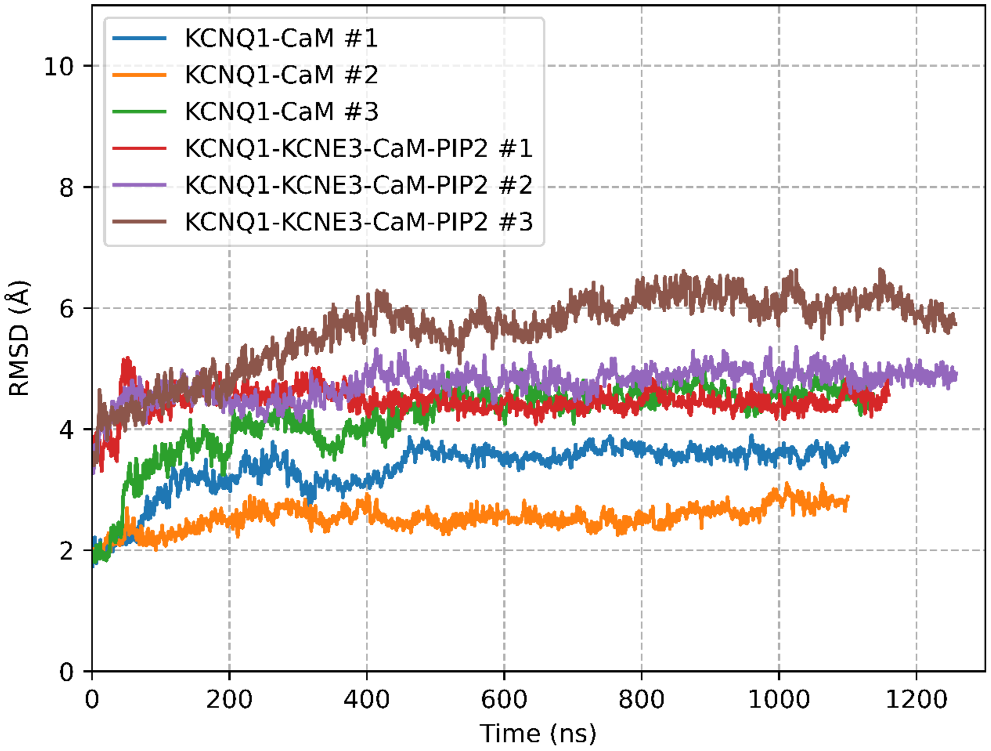


**Supplementary Fig. 1. Root Mean Square Displacement (RMSD) profiles computed from the MD simulations of the KCNQ1-CaM and KCNQ1-KCNE3-CaM-PIP2 systems**. The RMSD calculation has been carried out considering the backbone atoms of the proteins from the corresponding initial structures.

**Supplementary Tables**

**Supplementary Table 1.** Length of each MD simulation and reference point after which the systems reach the convergence monitored via RMSD calculations.

| System | # Replica | Total length of the simulation | Convergence |
|---|---|---|---|
| KCNQ1-CaM (PDB: 6UZZ) | 1 | 1100 ns | After 500 ns |
| | 2 | 1100 ns | After 500 ns |
| | 3 | 1140 ns | After 500 ns |
| KCNQ1-KCNE3-CaM-PIP2 (PDB: 6V01) | 1 | 1160 ns | After 400 ns |
| | 2 | 1258 ns | After 400 ns |
| | 3 | 1256 ns | After 700 ns |

**Supplementary Table 2.** Minimal path lengths ($d_{min}$) and frequency ($f$) of the allosteric paths identified by the network analysis. The intra-subunit paths couple the source to the sink of the same n subunit while the inter-subunit paths the source to the sink of the n and n+1 subunits.

| KCNQ1-CaM (PDB: 6UZZ) | | | | | |
|---|---|---|---|---|---|
| Path family | | Intra-subunit | | Inter-subunit | |
| | | $d_{min} \pm \sigma$ | $f$ (%) | $d_{min} \pm \sigma$ | $f$ (%) |
| Inactivation | S1 | $15.04 \pm 3.10$ | 38 | $14.94 \pm 2.96$ | 45 |
| | P-Helix | $16.18 \pm 2.72$ | 60 | $15.51 \pm 2.47$ | 55 |
| | S6 | $13.43 \pm 0.59$ | 2 | No path | 0 |
| CaM-SF | S0-S1 | $26.19 \pm 2.95$ | 28 | $26.37 \pm 3.48$ | 29 |
| | S0-H45 | $30.49 \pm 2.29$ | 1 | $28.28 \pm 1.39$ | 1 |
| | S2 | $27.82 \pm 2.41$ | 19 | $27.99 \pm 2.82$ | 18 |
| | S3 | $29.42 \pm 2.26$ | 21 | $28.05 \pm 2.14$ | 15 |
| | HA | $39.45 \pm 1.36$ | 5 | $38.64 \pm 1.92$ | 7 |
| | HB | $34.67 \pm 2.51$ | 26 | $35.87 \pm 3.16$ | 30 |
| KCNQ1-KCNE3-CaM-PIP2 (PDB: 6V01) | | | | | |
| Inactivation | P-Helix | $15.93 \pm 1.77$ | 29 | $18.62 \pm 2.63$ | 40 |
| | S6 | $16.89 \pm 2.15$ | 71 | $20.11 \pm 2.44$ | 60 |
| CaM-SF | HA | $34.52 \pm 3.32$ | 100 | $34.39 \pm 3.37$ | 100 |

**Supplementary Table 3**. **Betweenness of residues along the inactivation paths computed in the KCNQ1-CaM system.** The error is expressed as standard deviation.

| Residue | Betweenness | Standard deviation |
|---|---|---|
| S140 | 0.81 | 0.36 |
| V141 | 1.70 | 0.29 |
| R228 | 1.84 | 0.52 |
| I230 | 1.11 | 0.00 |
| R231 | 1.03 | 0.01 |
| I235 | 1.47 | 0.03 |
| L236 | 2.14 | 0.04 |
| L273 | 1.66 | 0.19 |
| I274 | 1.47 | 0.23 |
| F275 | 1.36 | 0.48 |
| S276 | 2.42 | 0.65 |
| S277 | 1.89 | 1.07 |
| A300 | 1.06 | 0.00 |
| D301 | 2.13 | 0.00 |
| W305 | 2.98 | 0.47 |
| V307 | 0.96 | 0.03 |
| V308 | 2.22 | 0.01 |
| I313 | 3.18 | 0.37 |
| Y315 | 3.73 | 0.31 |

**Supplementary Table 4**. **Betweenness of residues along the inactivation paths computed in the KCNQ1-KCNE3-CaM-PIP2 system.** The error is expressed as standard deviation.

| Residue | Betweenness | Standard deviation |
|---|---|---|
| R231 | 0.25 | 0.00 |
| F232 | 0.31 | 0.02 |
| Q234 | 0.38 | 0.00 |
| I235 | 0.35 | 0.00 |
| I274 | 1.16 | 0.15 |
| S277 | 1.25 | 0.00 |
| F279 | 1.20 | 0.55 |
| V280 | 1.13 | 0.19 |
| Y281 | 1.02 | 0.06 |
| G306 | 2.20 | 0.52 |
| T309 | 0.63 | 0.00 |
| I313 | 3.19 | 0.42 |
| Y315 | 4.63 | 0.25 |
| P320 | 1.50 | 0.10 |
| I328 | 1.25 | 0.00 |
| A329 | 0.11 | 0.00 |

**Supplementary Table 5. Betweenness of residues along the CaM-SF paths computed in the KCNQ1-CaM system.** The error is expressed as standard deviation.

| Residue | Betweenness | Standard deviation |
|---|---|---|
| R116 | 0.39 | 0.05 |
| P117 | 0.12 | 0.01 |
| H126 | 0.24 | 0.02 |
| V129 | 0.50 | 0.04 |
| V133 | 0.99 | 0.08 |
| L137 | 1.98 | 1.05 |
| R174 | 0.02 | 0.00 |
| S177 | 0.01 | 0.01 |
| C180 | 1.59 | 0.10 |
| R181 | 1.89 | 0.01 |
| S182 | 12.33 | 0.61 |
| Y184 | 2.17 | 0.56 |
| F193 | 1.15 | 0.83 |
| P197 | 0.29 | 0.03 |
| I201 | 2.70 | 1.47 |
| I235 | 2.71 | 1.20 |
| L236 | 2.28 | 0.32 |
| L239 | 7.12 | 1.70 |
| H240 | 0.45 | 0.07 |
| Q244 | 1.71 | 0.09 |
| G245 | 0.06 | 0.00 |
| T247 | 0.23 | 0.02 |
| Y267 | 2.82 | 0.63 |
| F270 | 3.00 | 0.58 |
| L273 | 1.75 | 1.21 |
| I274 | 3.31 | 0.58 |
| F275 | 2.84 | 0.13 |
| S276 | 1.73 | 0.47 |
| S277 | 1.88 | 0.20 |
| V280 | 1.50 | 0.91 |
| Y299 | 1.17 | 0.17 |
| A302 | 1.54 | 0.70 |
| W305 | 1.77 | 0.26 |
| G306 | 3.49 | 0.18 |
| T309 | 1.17 | 1.00 |
| T312 | 5.39 | 0.13 |
| I313 | 2.60 | 0.60 |
| Y315 | 4.59 | 0.12 |

| | | |
|---|---|---|
| G316 | 2.88 | 0.75 |
| V319 | 4.38 | 3.66 |
| P320 | 3.50 | 1.74 |
| I337 | 1.36 | 0.48 |
| S338 | 4.55 | 1.01 |
| L342 | 2.40 | 1.92 |
| F351 | 2.50 | 0.18 |
| K354 | 4.59 | 1.60 |
| Q357 | 4.96 | 0.17 |
| K358 | 2.93 | 0.69 |
| Q361 | 1.46 | 0.34 |
| H363 | 1.40 | 0.36 |
| R366 | 2.02 | 0.01 |
| Y522 | 0.30 | 0.00 |
| K526 | 0.53 | 0.20 |
| Q530 | 1.13 | 0.15 |
| K534 | 1.40 | 0.82 |
| P535 | 1.54 | 0.56 |

**Supplementary Table 6. Betweenness of residues along the CaM-SF paths computed in the KCNQ1-KCNE3-CaM-PIP2 system.** The error is expressed as standard deviation.

| Residue | Betweenness | Standard deviation |
|---|---|---|
| Q260 | 0.59 | 0.22 |
| T264 | 0.19 | 0.04 |
| T265 | 0.25 | 0.00 |
| T309 | 0.67 | 0.29 |
| V310 | 2.03 | 0.00 |
| Y315 | 7.44 | 0.59 |
| F340 | 2.50 | 0.24 |
| K358 | 0.17 | 0.00 |
| F364 | 0.60 | 0.02 |

**Supplementary Table 7. Betweenness of residues on CaM involved in the CaM-SF paths.** The error is expressed as standard deviation.

| Residue | Betweenness | Standard deviation |
|---|---|---|
| E46 | 0.15 | 0.15 |
| Q50 | 0.52 | 0.19 |
| N54 | 0.63 | 0.41 |
| A58 | 0.81 | 0.60 |
| M77 | 0.56 | 0.01 |
| E83 | 0.61 | 0.20 |
| I86 | 0.13 | 0.31 |
| A89 | 0.88 | 0.11 |
| V92 | 1.05 | 0.00 |
| G97 | 3.03 | 0.61 |
| N98 | 3.51 | 0.56 |
| G99 | 2.89 | 0.42 |
| Y100 | 2.92 | 0.13 |

**Supplementary Table 8. Fitted values of voltage dependence of activation and their deduced free energy of WT Kv7.1 and its mutants.** Values in parentheses correspond to the number of independent recorded cells. Recordings were performed in 1.8 mM external $Ca^{2+}$ solutions and in nominally 0 µM free $Ca^{2+}$ internal pipette solution.

| | $V_{50}$ (mV) | z | $\Delta G_0$ (kcal/mol) | $\Delta\Delta G_0$ corrected (kcal/mol) |
|---|---|---|---|---|
| WT Kv7.1 (43) | -25.0 ± 1.4 | 2.1 ± 0.1 | -1.3 ± 0.1 | |
| S140A (11) | -28.3 ± 3.4 | 1.5 ± 0.1 | -1.0 ± 0.2 | 3.2 ± 1.9 |
| V141A (31) | -34.9 ± 2.1 | 2.8 ± 0.1 | -2.3 ± 0.2 | -1.8 ± 0.3 |
| C180A (9) | -22.8 ± 1.1 | 3.2 ± 0.2 | -1.7 ± 0.3 | -1.4 ± 0.3 |
| R181A (11) | -28.1 ± 2.1 | 3.3 ± 0.2 | -2.1 ± 0.2 | -0.7 ± 0.2 |
| S182A (13) | -23.3 ± 2.3 | 3.3 ± 0.1 | -1.7 ± 0.1 | -4.5 ± 0.2 |
| K183A (9) | -19.7 ± 1.7 | 3.3 ± 0.3 | -1.5 ± 0.2 | -0.2 ± 0.2 |
| R228W (8) | 11.1 ± 2.9 | 1.2 ± 0.1 | 0.3 ± 0.1 | 9.1 ± 0.3 |
| I230W (8) | -8.8±2.4 | 1.8±0.2 | -0.3±0.1 | 2.4 ± 0.1 |
| L233W (12) | -35.7 ± 2.6 | 3.6 ± 0.5 | -2.9 ± 0.3 | -2.8 ± 0.2 |
| Q234W (6) | 2.2±4.0 | 1.3±0.1 | 0.1±0.1 | 2.5 ± 0.2 |
| I235A (13) | 33.5 ± 2.5 | 2.3 ± 0.1 | 1.8 ± 0.1 | 3.6 ± 0.2 |
| L236W (10) | 9.8±1.5 | 2.7±0.2 | 0.6±0.1 | 4.0 ± 0.3 |
| L239W (5) | 39.5 ± 3.6 | 2.4 ± 0.3 | 2.2 ± 0.2 | 6.1 ± 0.2 |
| L273A (6) | No expression | | | |
| L273F (11) | -41.6 ± 2.6 | 3.4 ± 0.6 | -3.3 ± 0.4 | -12.4 ± 0.4 |
| I274A (10) | -33.6 ± 3.6 | 2.5 ± 0.3 | -2.1 ± 0.4 | -0.9 ± 0.5 |
| G306A (6) | -32.1 ± 1.4 | 3.3 ± 0.4 | -2.4 ± 0.3 | -3.9 ± 1.0 |
| V307L (8) | -27.5 ± 2.0 | 4.4 ± 1.2 | -2.8 ± 0.4 | -5.3 ± 0.7 |
| V308A (10) | -5.7 ± 2.5 | 1.1 ± 0.0 | -0.2 ± 0.1 | 2.0 ± 0.1 |
| T312S (8) | -22.3 ± 7.6 | 2.3 ± 0.3 | -1.2 ± 0.3 | 0.3 ± 0.2 |
| S338F (8) | -21.5 ± 2.1 | 2.4 ± 0.3 | -1.2 ± 0.2 | 0.1 ± 0.1 |
| F351A (8) | 34.5 ± 3.2 | 2.7 ± 0.4 | 2.1 ± 0.3 | 3.4 ± 0.4 |
| K354W (7) | -3.5 ± 3.2 | 1.5 ± 0.3 | -1.4 ± 0.2 | -0.2 ± 0.2 |
| Q357W (13) | -10.2 ± 2.5 | 1.5 ± 0.2 | -0.4 ± 0.1 | 1.3 ± 0.3 |
| K358W (7) | -30.6 ± 11.0 | 1.8 ± 0.3 | -1.3 ± 0.4 | 0 |
| F364A (6) | -18.6 ± 5.5 | 1.5 ± 0.2 | -0.7 ± 0.3 | 0.6 ± 0.3 |
| R366W (11) | 8.8 ± 3.1 | 1.5 ± 0.6 | 0.3 ± 0.2 | 7.3 ± 1.2 |
| K526E | -18.3 ± 4.0 | 1.9 ± 0.4 | -0.8 ±0.3 | 1.3 ± 0.5 |
| Kv7.1 + Calmodulin | $V_{50}$ (mV) | z | $\Delta G_0$ (kcal/mol) | $\Delta\Delta G_0$ corrected (kcal/mol) |
| WT (10) | -25.1 ± 1.2 | 3.4 ± 0.5 | -2.0 ± 0.4 | |
| Y100A (12) | -23.5 ± 2.1 | 3.3 ± 0.3 | -1.8 ± 0.2 | 0.2 ± 0.2 |
| Y100D (12) | -16.5 ± 1.2 | 2.4 ± 0.4 | -0.9 ± 0.4 | 1.5 ± 0.5 |
| N138W (12) | -35.5 ± 2.2 | 4.4 ± 0.6 | -3.6 ± 0.8 | -1.6 ± 0.5 |
